# Optogenetic induction of appetitive and aversive taste memories in *Drosophila*

**DOI:** 10.1101/2021.11.12.468444

**Authors:** Meghan Jelen, Pierre-Yves Musso, Pierre Junca, Michael D. Gordon

**Affiliations:** Department of Zoology and Life Sciences Institute, University of British Columbia, Vancouver, Canada

## Abstract

Tastes are typically thought to evoke innate appetitive or aversive behaviours, prompting food acceptance or rejection. However, research in *Drosophila melanogaster* indicates that taste responses can be modified through experience-dependent changes in mushroom body circuits. In this study, we develop a novel taste learning paradigm using closed-loop optogenetics. We find that appetitive and aversive taste memories can be formed by pairing gustatory stimuli with optogenetic activation of sensory or dopaminergic neurons associated with reward or punishment. As with olfactory memories, distinct dopaminergic subpopulations drive the parallel formation of short- and long-term appetitive memories. Long-term memories are protein synthesis-dependent and have energetic requirements that are satisfied by a variety of caloric food sources or by direct stimulation of MB-MP1 dopaminergic neurons. Our paradigm affords new opportunities to probe plasticity mechanisms within the taste system and understand the extent to which taste responses depend on experience.

## INTRODUCTION

Food selection is influenced by a complex set of factors including external sensory input, interoceptive circuits signaling internal state, and plasticity driven by past feeding experiences. The gustatory system plays a critical role in evaluating the nutritional qualities of foods, and is generally thought to evoke innate appetitive or aversive behavioural responses. However, the degree to which taste processing can be modified by learning is unclear.

In flies, taste detection is mediated by gustatory receptor neurons (GRNs) located on the proboscis, pharynx, legs, wing margins, and ovipositor (Stocker, 1994). GRNs express a range of chemosensory receptors for detecting sugars, bitters, salts, and other contact chemical cues (Chen and Dahanukar, 2020). GRNs project to the subesophageal zone (SEZ) of the fly brain, where taste information is segregated based on modality, valence, and organ of detection (Marella et al., 2006; Thorne et al., 2004; Wang et al., 2004).

Although the valence of a specific taste is generally set, the intensity of the response can vary according to a variety of factors, including internal state. For example, starvation increases a fly’s sensitivity to sweet tastes and blunts bitter responses through direct modulation of GRN activity (Inagaki et al., 2014, 2012; LeDue et al., 2016; Marella et al., 2012). Moreover, flies lacking essential nutrients such as amino acids and salts exhibit increased nutrient-specific preference towards protein and salt-rich foods (Corrales-Carvajal et al., 2016; Jaeger et al., 2018; Steck et al., 2018).

In addition to internal state-dependent changes in nutrient drive, fly taste responses can be altered by experience. Most notably, short-term taste-specific suppression of appetitive responses can be achieved through pairing with either bitter taste or noxious heat (Keene and Masek, 2012; Kirkhart and Scott, 2015a; Masek et al., 2015; Tauber et al., 2017). This plasticity requires an integrative memory association area called the mushroom body (MB), which is known to represent stimuli of different modalities, including taste (Keene and Masek, 2012; Kirkhart and Scott, 2015a; Masek et al., 2015). Thus, while taste responses are carried out by innate circuits, they also exhibit experience-dependent changes driven by the adaptable networks of the MB (Colomb et al., 2009; Kirkhart and Scott, 2015a; Krashes et al., 2009).

The MB is composed of approximately ∼4,000 intrinsic Kenyon cells (KCs), whose dendrites receive inputs from different sensory systems (Kirkhart and Scott, 2015a; Schwaerzel et al., 2003; Tanaka et al., 2008; Vogt et al., 2014). KCs form *en passant* synapses with mushroom body output neurons (MBONs), and MBONS send projections to neuropils outside of the MB to modulate behavioural output (Crittenden et al., 1998; Tanaka et al., 2008). Heterogeneous dopaminergic neurons (DANs) project to specific MB compartments and are activated in response to rewarding or punishing stimuli (Burke and Waddell, 2011; Gervasi et al., 2010; Mao and Davis, 2009; Tomchik and Davis, 2009). Coincident KC activation and release of dopamine results in the depression of corresponding KC-MBON synapses and skews the MB network towards approach or avoidance behaviours (Aso et al., 2014; Cohn et al., 2015; Perisse et al., 2013).

Interestingly, direct activation of DANs can function as an Unconditioned Stimulus (US) in some *Drosophila* associative learning paradigms (Aso et al., 2012; Claridge-Chang et al., 2009; Colomb et al., 2009; Liu et al., 2012). Optogenetic or thermogenetic activation of the Protocerebral Anterior Medial (PAMs) neurons, a rewarding DAN subpopulation, following or in coincidence with an odor Conditioned Stimulus (CS) results in the formation of an appetitive memory, while activation of punishing PPL1 DANs leads to the formation of an aversive memory (Masek et al., 2015; Yamagata et al., 2015). Similarly, phasic optogenetic activation of specific dopaminergic subsets in the absence of a physical reward can lead to the formation of conditioned behaviours in mice (Saunders et al., 2018).

Appetitive Short- and Long-Term Memories (STM; LTM) are formed by independent PAM subpopulations, with the nutritional value of the sugar reward and satiation state of the fly contributing to the strength of the association (Burke and Waddell, 2011; Colomb et al., 2009; Musso et al., 2015; Yamagata et al., 2015). Whereas STM may be formed under more flexible conditions with a sweet tasting reward, the formation of LTM requires a nutritious sugar (Burke and Waddell, 2011; Musso et al., 2015). Caloric sugars gate memory consolidation by promoting sustained rhythmic activity of MB-MP1 DANs (Musso et al., 2015; Plaçais et al., 2017, 2012). Interestingly, this signaling may occur up to 5 hours post-ingestion, suggesting that there is a critical time window for the formation of LTM (Musso et al., 2015; Pavlowsky et al., 2018).

Although flies are known to exhibit aversive short-term taste memories, where appetitive taste responses are diminished through punishment (Kim et al., 2017; Kirkhart and Scott, 2015; Masek et al., 2015; Masek and Scott, 2010), the full extent to which taste behaviours are modifiable by learning is unknown. Can taste responses be enhanced by appetitive conditioning? Can flies form long-term memories about taste? These are difficult questions to answer using traditional methods for several reasons. First, appetitive association paradigms generally rely on food as the US, which could interfere with the representation of a taste CS and also modify future taste behaviours through changes in satiety state. Second, repeated temporal pairing of a taste CS with a US is difficult to achieve in flies without immobilization, making long-lasting memories difficult to test. To circumvent these issues, we developed an optogenetic learning paradigm that facilitates rapid, repeated CS/US pairing while maintaining similar satiation states between groups. In this paradigm, we couple a taste (the CS) with optogenetic GRN or dopaminergic neuron stimulation (the US) in order to study conditioned taste responses or ‘taste memories’. For the purpose of our study, we will define a taste memory as a measurable change in a fly’s behavioural response to a previously encountered taste stimulus.

Using our novel paradigm, we show that flies are capable of forming both appetitive and aversive short- and long-term taste memories. As in olfaction, appetitive taste memories are driven by discrete PAM populations, and activation of a single PAM subpopulation is sufficient to induce appetitive LTM. The formation of appetitive LTM requires *de novo* protein synthesis and is contingent on caloric intake. Moreover, sugar, certain amino acids, and lactic acid can provide the energy required to support LTM formation, and this requirement is also satisfied by thermogenetic activation of MB-MP1 neurons.

## RESULTS

### Pairing GRN activation with a food source leads to taste memory formation

We previously developed a system called the sip-triggered optogenetic behavioural enclosure (STROBE), in which individual flies are placed in an arena with free access to two odorless food sources (Musso et al., 2019). Interactions (mostly sips) with one food source triggers nearly instantaneous activation of a red LED, which can be used for optogenetic stimulation of neurons expressing CsChrimson. We reasoned that, if sipping on a tastant (the CS^+^) triggers activation of neurons that provide either positive or negative reinforcement, we may observe a change in the number of interactions a fly initiates upon subsequent exposure to the same CS^+^ (Figure 1A).

**Figure 1:**
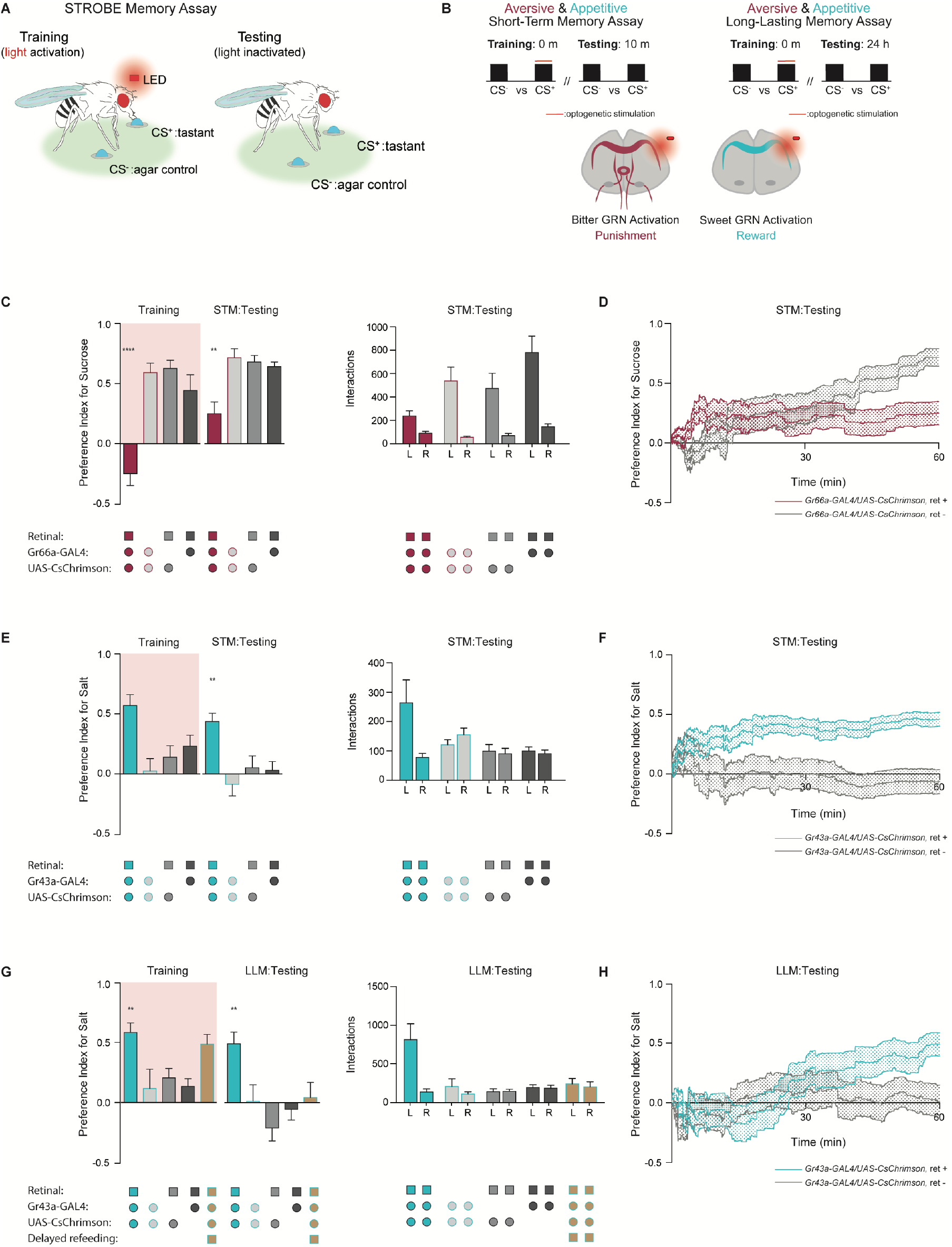
GRNs produce punishment and reward signals capable of facilitating taste memory formation. (A) Diagram outlining STROBE memory paradigm. <u>Training:</u> starved flies freely interact with a LED-activating tastant for 40 minutes. *CsChrimson* induces bitter or sweet neuron stimulation upon LED-activation, pairing feeding with punishment or reward. <u>Testing:</u> associative memory is measured by assessing preference for the tastant compared to agar for a 1-hour time period. (B) Schematic of aversive and appetitive STM and LLM timelines and a diagram of activated bitter and sweet neurons in the SEZ. (C) Preference indices (left) and tastant interaction numbers (right) for *Gr66a>CsChrimson* flies compared to genetic controls during training and 10 minutes later upon testing. (D) Cumulative average preference indices over the course of testing in (C), (n=16-30). (E) Preference index (left) and interactions numbers (right) for *Gr43a>CsChrimson* flies fed all-*trans-*retinal compared to controls in the short-term memory assay. (F) Preference index of flies in (E) throughout testing (n= 12-23). (G) Preference index (left) and interactions numbers (right) for *Gr43a>CsChrimson* flies fed all-*trans-*retinal compared to controls in the long-lasting memory assay. (H) Average preference index as a function of time for the testing period of flies in the long-lasting memory assay (n= 14-30). Preference index is mean ± SEM, One-way ANOVA, Dunnett’s post hoc test: \*\**p* < 0.01, \*\*\*\**p* < 0.0001.

We began by testing the efficacy of the STROBE in inducing aversive and appetitive memories through optogenetic activation of bitter and sweet GRNs, respectively (Figure 1B). Bitter GRN stimulation is known to activate PPL1 DANs, while sweet GRNs activate PAMs (Keene and Masek, 2012; Kirkhart and Scott, 2015; Liu et al., 2012; Masek et al., 2015). Moreover, bitter or sweet GRN activation with *Gr66a-* or *Gr43a-Gal4* is sufficient for STM induction in taste and olfactory associative learning paradigms (Keene and Masek, 2012; Yamagata et al., 2015). Therefore, we tested whether pairing GRN activation with feeding on a single taste modality could create an associative taste memory that altered subsequent behaviour to the taste.

In the aversive taste memory paradigm, interactions with 25 mM sucrose (CS^+^) during training triggered LED activation of Gr66a bitter neurons expressing CsChrimson (Figure S1A), which led to CS^+^ avoidance relative to plain agar (CS^-^) (Figure 1C). During testing, we disable the STROBE lights and measured preference towards 25 mM sucrose (CS^+^) relative to agar (CS^-^) to see if flies have formed aversive taste memories. Indeed, ten minutes after training, flies that experienced bitter GRN activation during training showed a lower sugar preference than control flies of the same genotype that were not fed the obligate CsChrimson cofactor all-*trans*-retinal, and retinal-fed control genotypes carrying either the Gal4 or UAS alone. This preference change reflected a non-significant trend towards decreased interactions with the CS^+^ compared to controls (Figure 1C). Examining the preference indices over time revealed that both trained flies and controls exhibited similar sugar preference after the first 30 minutes of testing, but the difference between groups emerged from a steady increase in sugar preference in the control group while trained flies’ preference remained stable (Figure 1D). A similar effect was produced by the activation of PPK23^glut^ ‘high salt’ GRNs, which also carries a negative valence in salt-satiated flies (Figure S1A, C). Importantly, these effects are not due to heightened satiety in trained flies because training in this paradigm is associated with fewer food interactions than controls.

For appetitive training, we chose 75 mM NaCl as the CS^+^, since flies show neither strong attraction nor aversion to this concentration of salt (Zhang et al., 2013). Interactions with the CS^+^ in this paradigm triggered optogenetic activation of sweet neurons, either with *Gr43a-Gal4*, which labels a subset of leg and pharyngeal sweet neurons in addition to fructose-sensitive neurons in the protocerebrum, or *Gr64f-Gal4*, which labels most peripheral sweet GRNs (Figure S1A). In both cases, sweet GRN activation produced an increased preference for the salt CS^+^ during training and also testing 10-minutes later. (Figure 1E and Figure S1C). The increased preference is evident early during testing and maintained throughout the testing phase (Figure 1F). Like the aversive memory paradigm, the effects of appetitive conditioning cannot easily be explained through changes in internal state, since trained flies interacted more with the food during training and therefore should have a lower salt drive during testing. Interestingly, refeeding flies with standard medium directly after training in the appetitive paradigm led to a long-lasting preference for the CS^+^, revealed by testing 24-hours later (Figure 1G, 1H and Figure S1E). This stands in contrast to the aversive paradigm, where reduced preference for sugar following bitter GRN activation was absent 24 hours later (Figure S1A, B).

### DAN activation is sufficient for the induction of short and long-lasting taste memories

We next asked whether direct activation of DANs during feeding could drive the formation of taste memories. Aversive short-term taste memory depends on multiple PPL1 DANs, including PPL1-α’2 α2 and PPL1-α3 (Masek et al., 2015), while appetitive short-term taste memories have not been previously reported. We first tested whether activating PPL1 DANs coincident with tastant interactions would lead to STM formation in the STROBE. Stimulation of PPL1 neurons reduced sucrose preference during training, and a reduced preference was also observed during short-term memory testing 10 minutes later (Figure 2A). This decreased preference was sustained throughout the entire period of testing (Figure 2B). Interestingly, unlike activation of bitter sensory neurons, PPL1 activation also produced a long-lasting aversive memory that was expressed 24 hours after training and remained stable through the duration of testing (Figure 2C, D).

**Figure 2:**
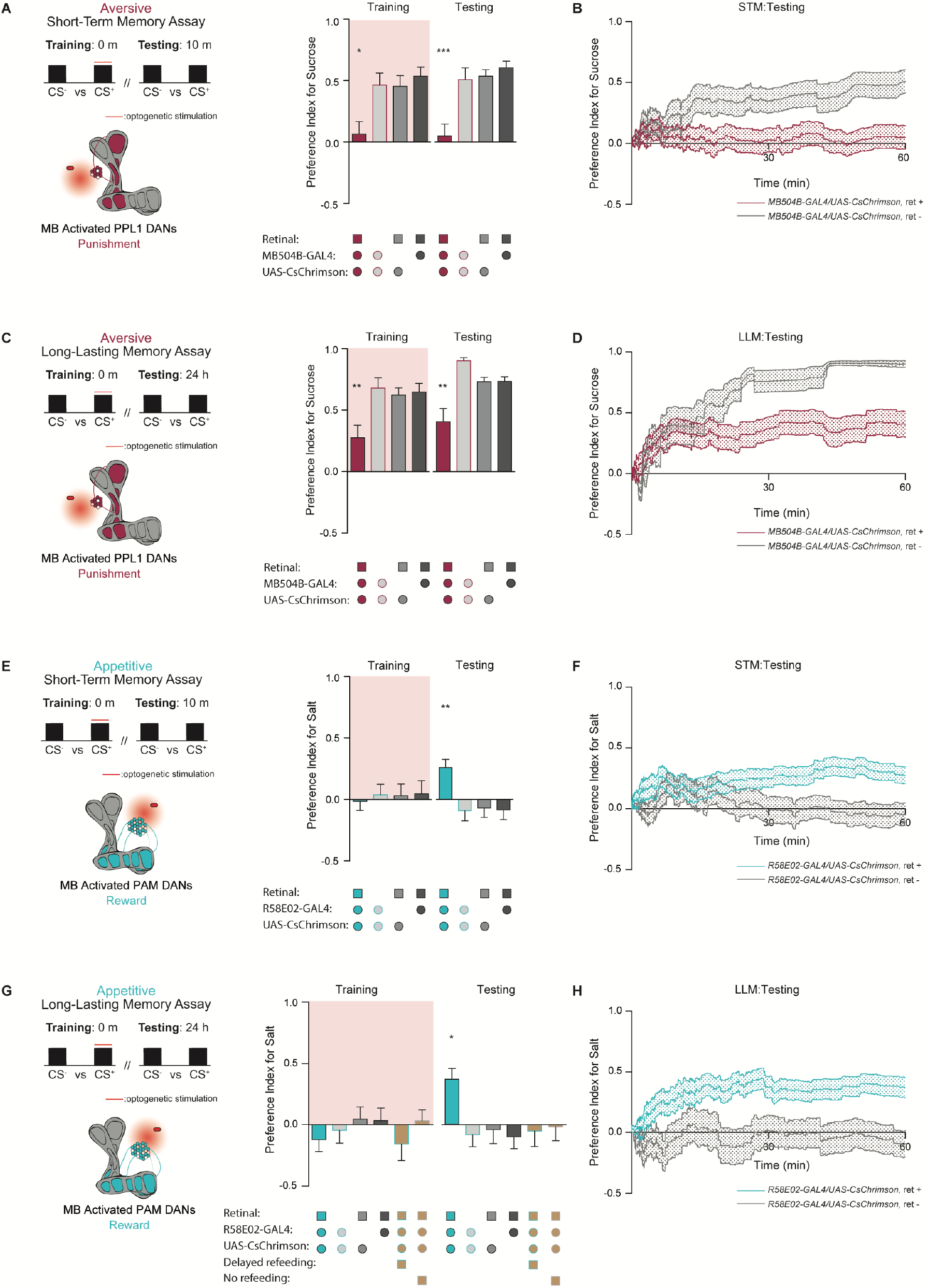
PPL1 and PAM neural activation is sufficient for the induction of short and long-lasting taste memories. (A, B) Paradigm timeline schematic and MB model indicating PPL1 compartments activated by optogenetics. Preference indices for *MB504B>CsChrimson* flies and controls as an average (A), and over time (B) in the short-term memory assay (n=19-31). (C, D) Schematic for the long-lasting memory assay, and preference index for *MB504B>CsChrimson* flies compared to genetic controls as an average (C), and corresponding time curve (D) for the duration of testing (n=20-33). (E, F) Assay timeline, and MB model with activated PAM compartments highlighted. Preference index for *R58E02>CsChrimson* flies comparted to controls (E) and as a function of time (F) in the short-term memory assay (n=25-38). (G, H) Preference index for *R58E02>CsChrimson* flies compared to controls (E) and over the total duration of testing (F) in the long-lasting memory assay (n=17-35). Preference indices are mean ± SEM, One-way ANOVA, Dunnett’s post hoc test: \**p* < 0.05, \*\**p* < 0.01, \*\*\**p* < 0.001.

To test the effect of appetitive DAN activation, we used flies expressing CsChrimson in PAM neurons under control of *R58E02-Gal4*. Intriguingly, although optogenetic activation of PAM neurons signals reward to the MB, it did not affect preference towards light-paired 75 mM NaCl (CS^+^) during training. Nonetheless, this pairing resulted in appetitive memory expression during testing 10 minutes and 24 hours after training (Figure 2E, G). These taste memories were stable throughout the entire duration of testing (Figure 2F, H). Thus, optogenetic activation of PAM neurons in the STROBE was able to write both short- and long-lasting appetitive taste memories in the absence of acute effects on feeding. Importantly, flies are also able to form appetitive memories to an alternative CS^+^ tastant, monopotassium glutamate (MPG) (Figure S2A). Taste memories were specific for CS^+^ taste identity, as flies trained with NaCl as the CS^+^ and MPG as the CS^-^ showed a clear preference for NaCl during testing (Figure S2B). Moreover, flies trained with NaCl as the CS^+^ do not show an elevated preference to MPG when it is introduced as a novel tastant during testing (Figure S2C). In addition to demonstrating specificity for the CS^+^, these experiments show that the observed change in behavior is not a memory of food position within the arena.

Consistent with long-term olfactory memories induced by DAN activation, we found long-lasting taste memory required an energetic food source, and therefore flies were refed for a brief period after training (Figure 2G). Flies that were fed 7 hours post-training, after the memory consolidation time period defined in olfactory memory, did not express taste memories during testing (Figure 2G) (Musso et al., 2015). Thus, the contingencies governing the formation and expression of taste memories in *Drosophila* seem to be similar to those previously discovered for olfaction.

### The MB is required for the formation of short- and long-lasting taste memories

Prior research indicates that the intrinsic neurons of the MB are required for aversive taste memory formation (Masek et al., 2015). To confirm that the MB is required for appetitive taste memory formation, we silenced this neuropil throughout both our short-term and long-lasting memory assays using tetanus toxin expressed under control of the pan-MB driver *R13F02-LexA*. After pairing Gr43a activation with NaCl feeding, flies with silenced MBs did not exhibit elevated preference for salt during testing 10 minutes or 24 hours later. (Figure 3A, B). Similarly, PAM activation during feeding led to a sustained increase in preference for the NaCl tastant in control groups for both the STM and long-lasting memory assay, but not in flies with silenced MBs (Figure 3C, D). These findings indicate that MB intrinsic neurons play a pivotal role in the formation of appetitive taste memories (Figure 3F).

**Figure 3:**
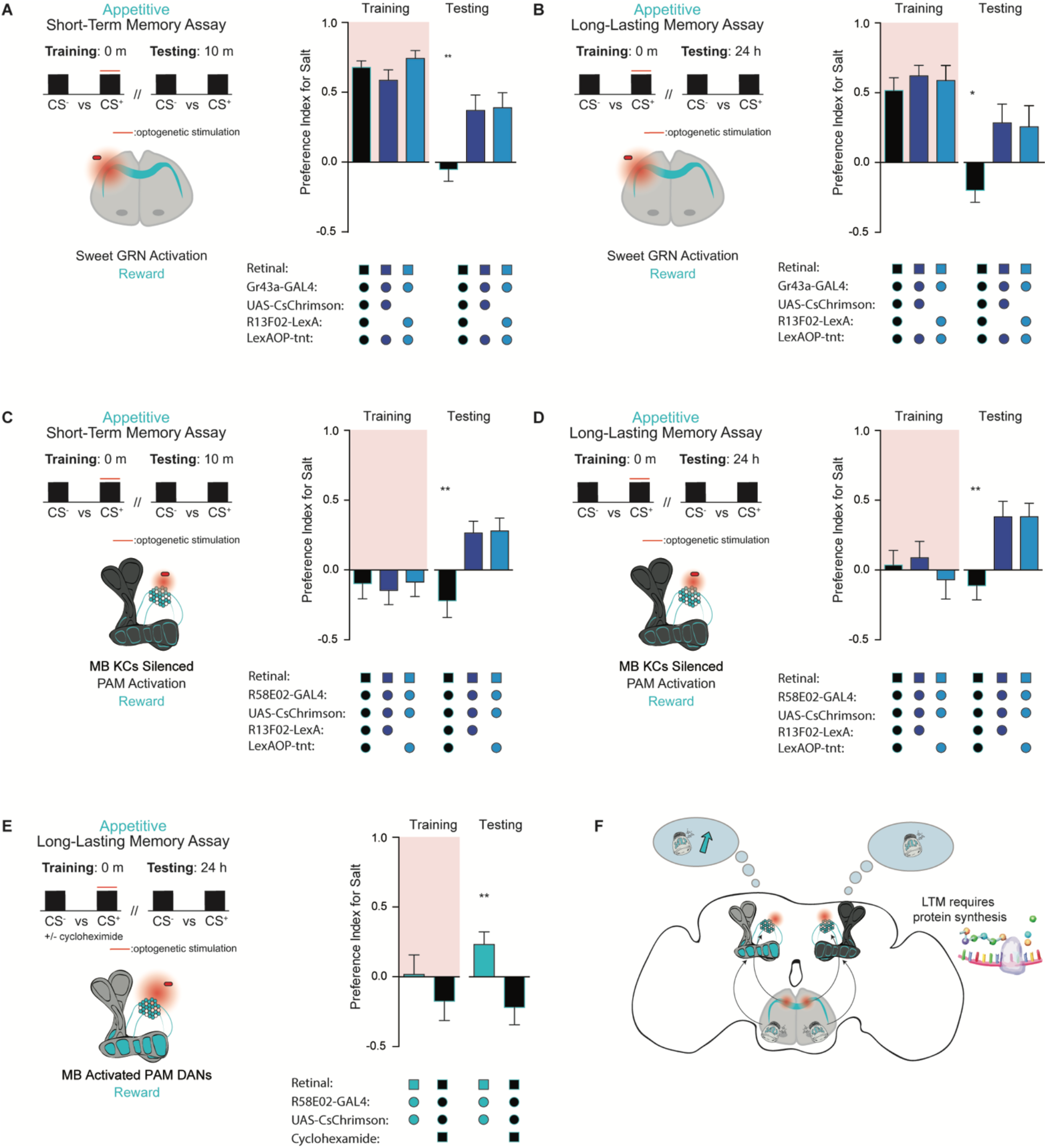
The MB is required for the formation of short- and long-term taste memories. (A, B) Preference indices for *Gr43a>CsChrimson* flies in the short-term (n=16-34) (A) and long-lasting (n=13-27) (B) memory assays when the MB is silenced, compared to controls. (C, D) Preference indices for *R58E02>CsChrimson* flies when the MB is silenced in the short-term (n=24-28)(C) and long-lasting (n=17-23) (D) memory paradigms, compared to controls. (E) Preference indices in the long-lasting memory assay for *R58E02>CsChrimson* flies fed protein synthesis inhibitor cycloheximide compared to vehicle-fed controls (n=17-22). (F) Model of appetitive taste memory formation via GRN/PAM activation. Preference indices are mean ± SEM, *t*-test/One-way ANOVA, Dunnett’s post hoc test: \**p* < 0.05, \*\**p* < 0.01.

To assess whether the molecular underpinnings of 24-hour appetitive taste memory are consistent with classic olfactory LTM, which requires *de novo* protein synthesis during memory consolidation, we fed flies all-*trans-*retinal laced with the protein synthesis inhibitor cycloheximide (CXM) (Colomb et al., 2009). As expected, flies fed CXM prior to training were unable to form long-term taste memories, in contrast to vehicle controls (Figure 3E). These results confirm that the taste memories being formed are protein synthesis dependent, and can be considered long-term memories (Figure 3F).

### Distinct PAM subpopulations induce appetitive short- and long-term taste memories

Distinct, non-overlapping subpopulations of PAM neurons, labeled by *R48B04-Gal4* and *R15A04-Gal4*, mediate the formation of appetitive short- and long-term olfactory memories, respectively (Yamagata et al., 2015). Moreover, it has been hypothesized that two differential reinforcing effects of sugar reward – sweet taste and nutrition, are encoded by these segregated STM and LTM neural populations (Yamagata et al., 2015). We tested both populations in our appetitive STROBE memory assays to determine if the activation of these separate PAM clusters would support the formation of parallel short- and long-term taste memories. *R48B04>CsChrimson* flies formed appetitive short-term but not long-term taste memories, as shown by the higher salt preference of flies expressing active CsChrimson during STM testing but not LTM testing (Figure 4A, B). Conversely, activation of *R15A04-Gal4* neurons produced LTM but not STM (Figure 4C, D). These results indicate that, much like appetitive olfactory memory, short- and long-term taste memories are formed in parallel by discrete PAM sub-populations.

**Figure 4:**
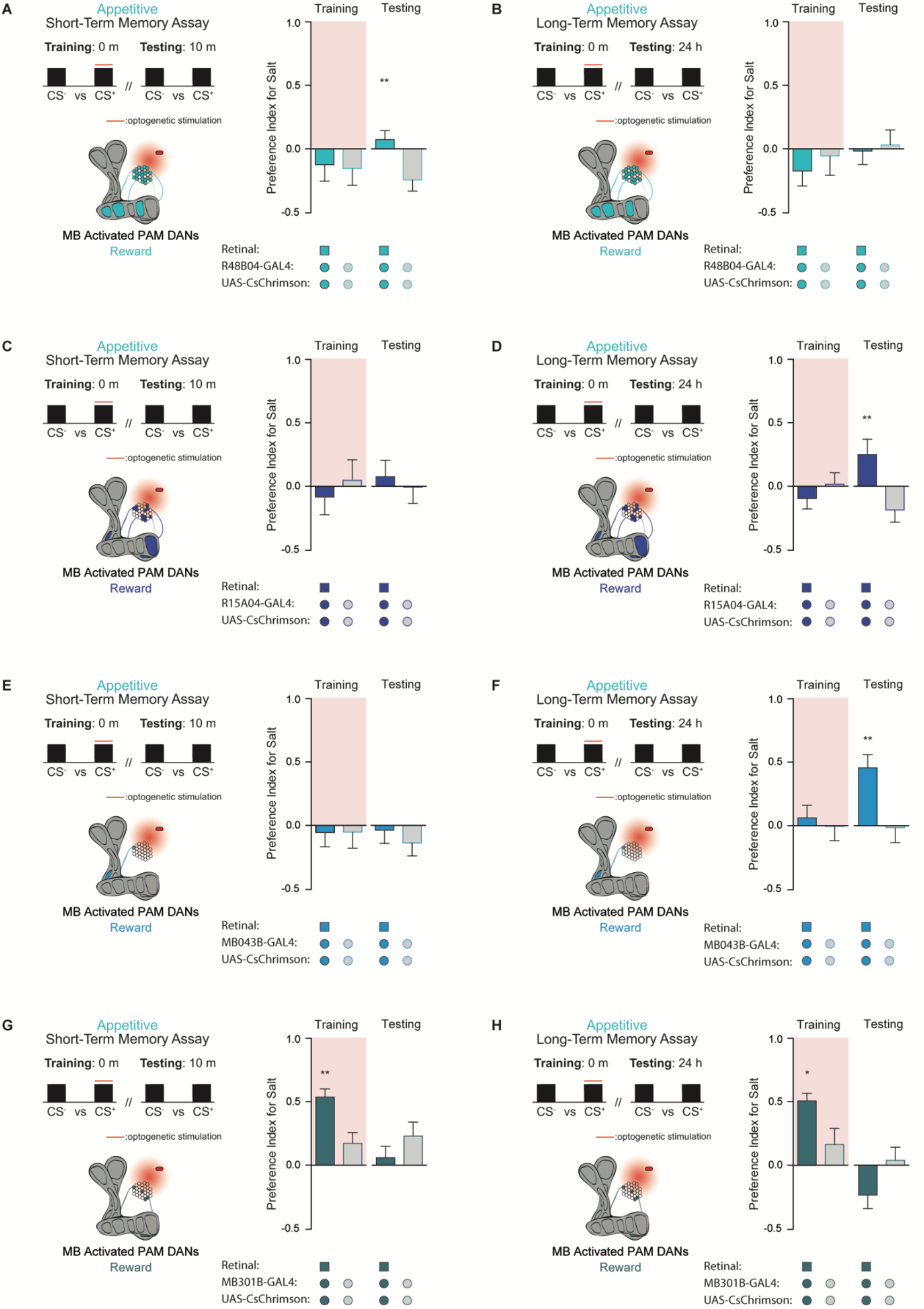
Discrete non-overlapping PAM subpopulations induce appetitive short- and long-term taste memories. (A, B) PAM subpopulation R48B04 innervates highlighted MB compartments (left), and preference indices of *R48B04>CsChrimson* flies for 75mM NaCl is tested in the short-term (n=21-28) (A), and long-term (n=15-17) (B) memory assays with or without retinal. (C, D). PAM subpopulation R15A04 innervates non-overlapping MB sub-regions compared to R48B04. Preference indices for *R15A04>CsChrimson* flies fed all-*trans*-retinal in the short-term (n=11-15) (C) and long-term (n=20-27) (D) taste memory assays with or without retinal. (E, F) PAM-α1 innervates a single compartment in the MB. Preference indices of *MB043B>CsChrimson* flies in the short-term (n=11-14) (E) and long-term (n=19-22) (F) memory assays with or without retinal. (G, H) PAM-β2β′2a synapses on the highlighted MB compartment. Preference indices for *MB301B>CsChrimson* flies during the short-term (n=20-27) (G) and long-term (n=10-15) (H) memory assays with or without retinal. Preference indices are mean ± SEM, *t*-test: \*\**p* < 0.01.

Next, we wondered whether activation of a single PAM cell subtype, PAM-α1, would be sufficient to induce taste memories. PAM-α1 neurons project to an MB compartment innervated by MBON-α1, which in turn feeds back onto PAM-α1 to form a recurrent reward loop necessary for the formation of appetitive olfactory LTM (Aso and Rubin, 2016; Ichinose et al., 2015). Consistent with its role in olfactory memory, activation of this PAM cell type in the STROBE with drivers *MB043B-Gal4* or *MB299B-Gal4* was sufficient to drive appetitive long-term, but not short-term, taste memory formation (Figure 4E, F and Figure S3A, B).

Interestingly, activation of another PAM subset labelled by *MB301B-Gal4* produced a higher preference for the salt CS during training, yet no sustained changes in taste preference during short- or long-term memory testing (Figure 4G, H). This demonstrates that the reward signaling associated with PAM cell activation occurs on multiple timescales to produce acute, short-, or long-term changes in behavior. Notably, the trend toward lower salt preference during testing in this experiment may reflect reduced salt drive due to increased salt consumption during training.

### Caloric food sources are required for the formation of associative long-term taste memories

Because refeeding with standard fly medium shortly after training is permissive for the consolidation of appetitive long-term taste memories, we next asked what types of nutrients would support memory formation. As expected, refeeding with L-glucose, a non-caloric sugar, did not lead to the formation of associative long-term taste memories (Figure 5A, B). However, along with sucrose, refeeding with lactic acid, yeast extract, and L-alanine promoted long-term memory, while L-aspartic acid did not. These results indicate that in addition to sucrose, other caloric nutrients can provide sufficient energy for long-term taste memory formation. Moreover, 7-hour delayed refeeding of each nutrient failed to support memory formation (Figure 5B). Thus, similar to olfactory LTM, the formation of appetitive taste LTM is dependent on an energy source being readily available during the memory consolidation window (Fujita and Tanimura, 2011; Musso et al., 2015).

**Figure 5:**
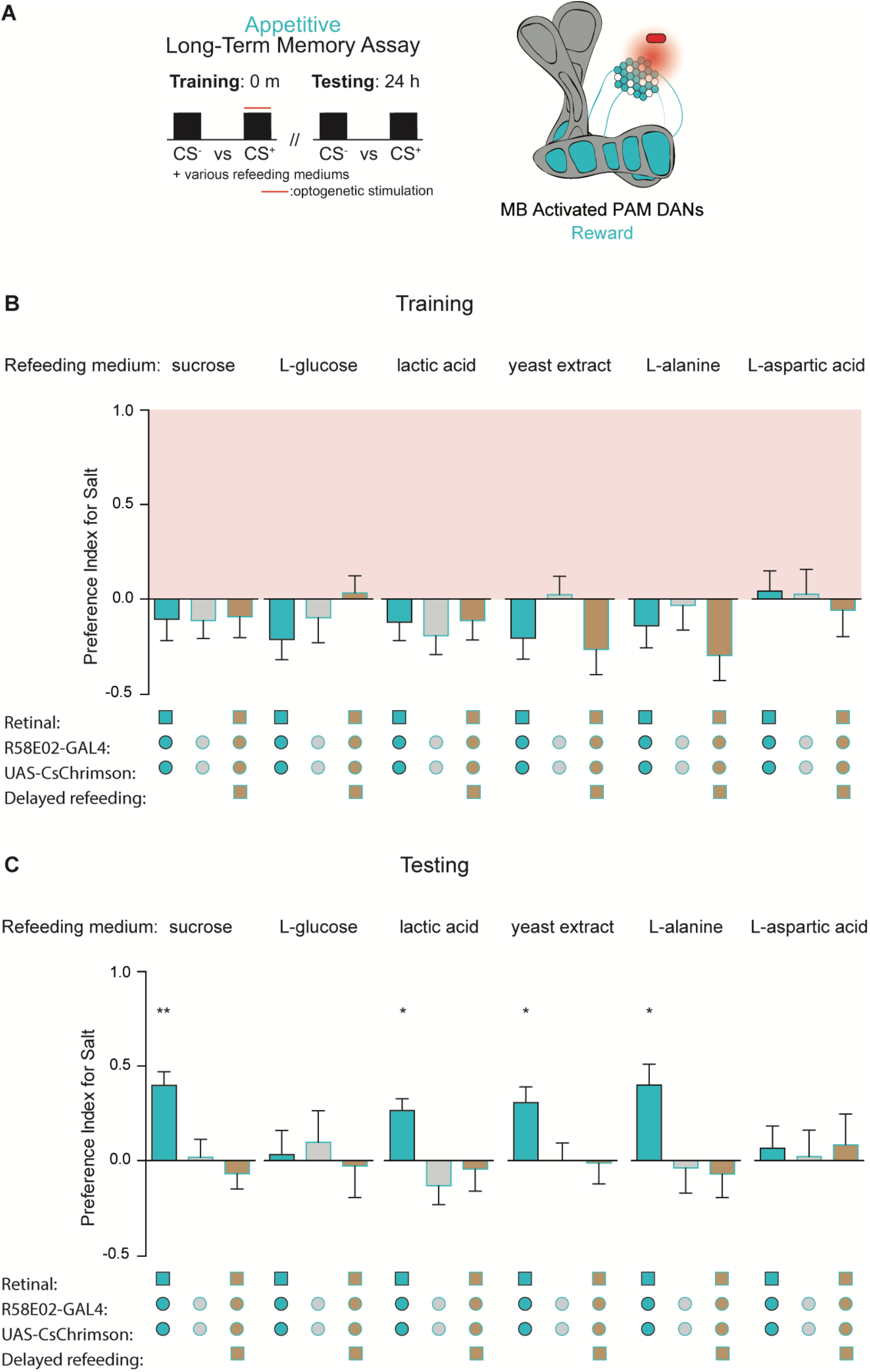
Caloric food sources are required for the formation of associative long-term taste memories. (A) Graphic of the timeline followed for the long-term taste memory paradigm, and the MB compartments innervated by PAM driver *R58E02-Gal4*. (B, C) Preference indices for *R58E02>CsChrimson* and control flies during training (B), and testing (C), after being refed with a caloric or non-caloric medium (n=13-28). Preference indices are mean ± SEM, One-way ANOVA, Dunnett’s post hoc test: \**p* < 0.05, \*\**p* < 0.01.

Our findings concerning the formation and expression of appetitive taste LTM bear striking similarities to those of olfactory LTM in terms of MB circuitry, dependence on protein synthesis, and energetic requirements. This led us to wonder if the energy gating performed by MB-MP1 neurons, which signal onto the mushroom body and promote energy flux in MB neurons during LTM, perform a similar function in taste memory (Musso et al., 2015; Plaçais et al., 2017, 2012). To test this hypothesis, we activated MB-MP1 neurons directly after training using *UAS-TRPA1* and delayed refeeding to outside the memory consolidation window. Compared to genetic controls, flies in which MB-MP1 neurons were activated post training showed significantly elevated memory scores during testing (Figure 6A, B). This confirms that MB-MP1 activation is sufficient to drive memory consolidation during long-term appetitive taste memory formation (Figure 6C).

**Figure 6:**
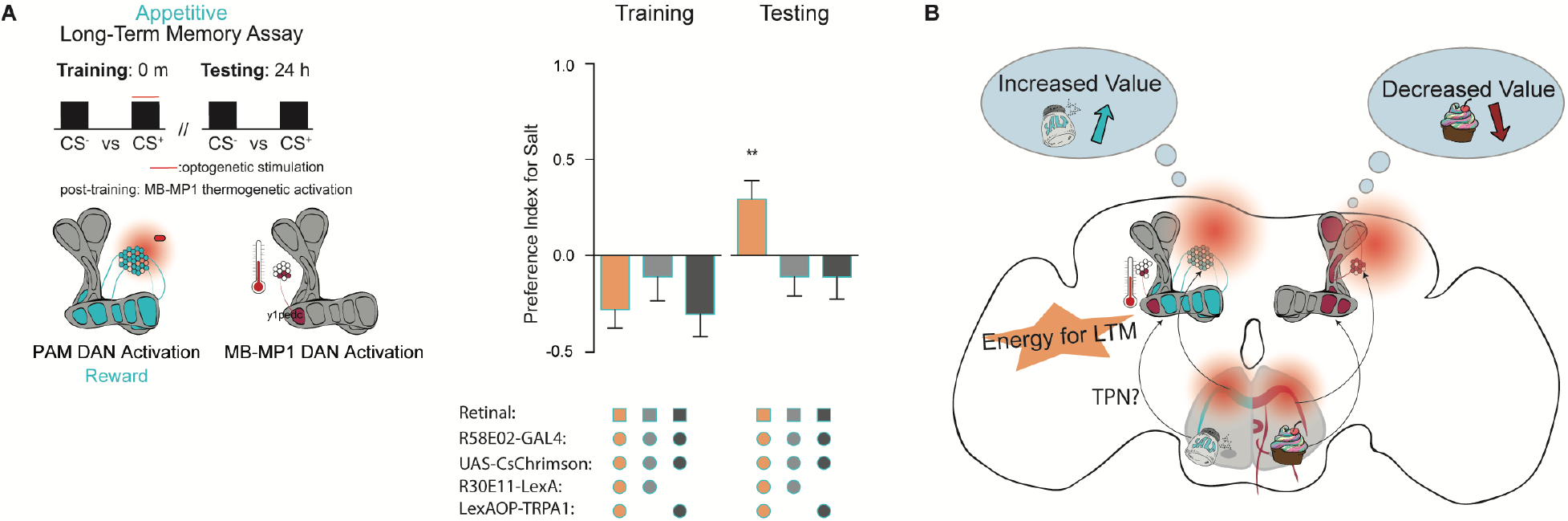
MB-MP1 neuron activation post-training replaces energy signal required for the formation of LTM. (A) Graphic of timeline followed for the LTM taste assay with thermogenetic activation of MB-MP1 neurons. Preference indices during training and testing for *R58E02>CsChrimson* flies fed all-*trans-*retinal, with MB-MP1 neurons thermogenetically activated post training using *R30E11>TRPA1*, compared to controls without MB-MP1 activation (n=18-29). (C) Schematic depicting summary of research. Preference indices are mean ± SEM, One-way ANOVA, Dunnett’s post hoc test: \*\**p* < 0.01.

## DISCUSSION

Gustation plays a vital role in determining the suitability of foods for ingestion. Yet, little is known about how experience influences higher-order taste representations and contributes to the continuous refinement of food selection. In fact, a memory system for the recollection of appetitive taste memories has not been described in flies. In this study, we use the STROBE to establish a novel learning paradigm and further investigate the formation and expression of taste memories. We demonstrate that flies are capable of forming short- and long-term appetitive and aversive taste memories towards two key nutrients – salt, and sugar. Much like olfactory memory, associative taste memory formation occurs within the MB and follows many of the same circuit and energetic principles.

We observed both positive and negative taste memories following optogenetic GRN stimulation concurrent with taste detection. The activation of bitter GRNs paired with sucrose led to the formation of STM, which agrees with previous research demonstrating that thermogenetic stimulation of bitter GRNs can act as a negative US, and lead to taste learning in short-term paradigms (Keene and Masek, 2012). Notably, although activation of sweet GRNs prompted both STM and LTM formation, bitter GRN activation was not sufficient for the formation of LTM in our assay. One possible explanation is that because bitter GRN activation strongly inhibits feeding, the number of pairings between sugar taste and bitter activation was insufficient to induce LTM. Consistent with this idea, PPL1 activation, which induced LTM, is less aversive during training and therefore allows more associations.

Although aversive taste memories have been established, evidence for appetitive taste memories has been sparse. Rats’ hedonic response to bitter compounds can be made more positive through pairing with sugar, and human studies suggest that children’s taste palates are malleable based on positive experiences with bitter vegetables (Breslin et al., 1990; Figueroa et al., 2020; Forestell and LoLordo, 2000; Wadhera et al., 2015). Thus, despite the difficulties of measuring taste memories in the lab, appetitive taste plasticity is very likely an ethologically important process.

The enhancement of salt palatability following co-incident activation of sweet taste may be surprising, given that NaCl on its own activates sweet GRNs (Jaeger et al., 2018; Marella et al., 2006). However, salt activates only about one third of sweet GRNs (Dweck et al., 2022), and thus appetitive memory formation may be driven by strong activation of the broader sweet neuron population. Nevertheless, directly stimulating DANs as the flies experienced taste inputs through feeding afforded us the ability to reduce this complication and also interrogate the roles of specific DAN populations. Taking a hypothesis driven approach, we confirmed that PAM neural subpopulations reinforce taste percepts much like olfactory inputs, and that short- and long-term memories are processed by distinct subpopulations. For example, activating β’2, γ4, and γ5 compartments with *R48B04-Gal4* produces STM in both olfactory and taste paradigms, while activation of α1, β’1, β2, and γ5 with *R15A04-Gal4* produces LTM in both. These results confirm that appetitive STM and LTM are processed in parallel in the MB (Trannoy et al., 2011; Yamagata et al., 2015). Given that tastes, like odours, activate the KC calyces (Kirkhart and Scott, 2015a), we speculate that optogenetic stimulation of PAM neurons during feeding modulates the strength of KC-MBON synaptic connections. Notably, activation of single PAM cell type produced different forms of memory in the STROBE. For example, stimulating PAM-α1 neurons during feeding drives appetitive taste LTM, while activation of PAM-β’1 was immediately rewarding.

A unique aspect of our long-term taste learning paradigm is that we uncoupled the US from a caloric food source. By doing this we were able to probe the energetic constraints gating LTM formation. For years it has been reported that long-term memory formation in *Drosophila* requires the intake of caloric sugar. Here, we demonstrate that the caloric requirements of long-term memory formation can be quenched by food sources other than sucrose, including lactic acid and yeast extract. Moreover, it seems that at least one amino-acid, L-alanine, is able to provide adequate energy, while L-aspartic acid cannot. We theorize that these foods may provide flies readily accessible energy, as neurons are able to metabolize both lactic acid and L-alanine into pyruvate to fuel the production of ATP via oxidative phosphorylation (de Tredern et al., 2021).

Energy gating in the MB is thought to be regulated by the MB-MP1-DANs. MB-MP1 neuron oscillations activate increased mitochondrial energy flux within the KCs, which is both necessary and sufficient to support LTM (Plaçais et al., 2017). To demonstrate sufficiency in our assay, we activated MB-MP1 neurons with TRPA1 directly after fly training, which effectively substitutes for a caloric food source and allows LTM formation (Figure 6C). These results suggest that MB-MP1 neurons integrate energy signals during formation of multiple types of LTM and may be influenced by a variety of caloric foods.

Overall, our results suggest that lasting changes in the value of specific tastes can occur in response to temporal association with appetitive or aversive stimuli, raising the possibility that such plasticity plays an important role in animals’ ongoing taste responses. Future experiments using the STROBE paradigm could further probe the molecular and circuit mechanisms underlying taste memories and advance our understanding of how taste preferences may be shaped by experience over an animal’s lifetime.

## MATERIALS AND METHODS

### Fly strains

Fly stocks were raised on a standard cornmeal diet at 25°C, 70% relative humidity. For neuronal activation, *20XUAS-IVS-CsChrimson*.*mVenus* (BDCS, stock number: 55135) was used. Dopaminergic PAM expression was targeted using previously described lines: *R58E02-GAL4* (Musso et al., 2015); *R58E02-LexA, R48B04-GAL4, R15A04-GAL4, R13F02-LexA*, and *R30E11-LexA* obtained from Bloomington (BDCS, stock numbers: 52740, 50347, 48671, 52460, 54209); and MB split-GAL4 lines *MB043B-GAL4, MB504B-GAL4, MB299B-GAL4, MB301B-GAL4* from Janelia Research Campus (Aso et al., 2014). GRN expression was driven using *Gr43a-GAL4, Gr64f-GAL4* (Dahanukar et al., 2007), *Gr66a-GAL4* (Wang et al., 2004), and *PPK23*^*glut*^*-GAL4* (*PPK23-GAL4, Gr66a-LexA::VP16, LexAop-Gal80* (Jaeger et al., 2018). *LexAop-tnt* was previously described (Liu et al., 2016). For temperature activation experiments *LexAop-TrpA1* was used (Liu et al., 2012).

### STROBE experiments

Mated female *Drosophila* were collected 2-3 days post eclosion and transferred into vials containing 1 ml of standard cornmeal medium supplemented with 1 mM all-*trans*-retinal (Sigma #R2500) or an ethanol vehicle control. Flies were maintained on this diet for 2 days in a dark environment. 24 hours prior to experimentation flies were starved at 25°C, 70% relative humidity, on 1% agar supplemented with 1 mM all-*trans*-retinal or ethanol vehicle control.

### STROBE training protocol

During the training phase for the short-term memory experiments the STROBE was loaded with 4 µL of tastant (salt: Sigma #S7653 or sucrose: Sigma #S7903) on channel 1 and 4 µL 1% agar on channel 2. The red LED was triggered only when a fly interacted with the tastant in channel 1. The duration of the training period was 40 minutes. For the STM training protocol, flies were then transferred to clean empty vials for 10 minutes while the experimental apparatus was cleaned. The training and testing phases of LTM experiments were performed as described for the STM experiments with the following exception: after the 40-minute training period flies, were transferred individually into vials containing standard cornmeal diet or nutrient of interest (sucrose: Sigma #S7903, L-glucose: Sigma #G5500, lactic acid: Sigma #69785, yeast extract: Sigma #Y1625, L-alanine: Sigma #05129, L-aspartic acid: Sigma #11230) and allowed to feed for 1 hour. They were then transferred into 1% agar starvation vials and kept at 18°C until the testing component of the experiment. For MB-MB1 activation experiments, after training flies were placed at 29°C, 70% relative humidity for 1 hour on 1% agar starvation vials. They were then transferred to 18°C and refed 8 hours later, outside of the memory consolidation. After 1 hour of feeding they were once again transferred into 1% agar starvation vials and kept at 18°C until the retrieval component of the experiment. The preference index for each individual fly was calculated as: (sips from channel 1 – sips from channel 2)/(sips from channel 1 + sips from channel 2). All experiments were performed with a light intensity of 11.2mW/cm^2^ at 25°C, 70% relative humidity.

### STROBE testing protocol

During testing, 4 µL of the same tastant (salt: Sigma #S7653, sucrose: Sigma #S7903, MPG: Sigma #G1501) was reloaded into channel one and 4 µL of 1% agar on channel 2. The optogenetic component of the system was deactivated such the red LED would no longer trigger if a fly interacted with the tastant. Flies were reloaded individually into the same arenas. The duration of the testing phase was 1 hour. The preference index for each individual fly was calculated as: (sips from channel 1 – sips from channel 2)/(sips from channel 1 + sips from channel 2).

### Immunofluorescence microscopy

Brain staining protocols were performed as previously described (Chu et al., 2014). Briefly, brains were fixed for 1 hour in 4% paraformaldehyde and dissected in PBS + 0.1% TritonX. After dissection brains were blocked in 5% NGS diluted with PBST for 1 hour. Brains were probed overnight at 4°C using the following primary antibody dilutions: rabbit anti-GFP (1:1000, Invitrogen #A11122), and mouse anti-brp (1:50, DSHB #nc82). After a 1hour wash period secondary antibodies goat anti-rabbit Alexa-488 (1:200, Invitrogen #A11008) and goat anti-mouse Alexa-568 (1:200, Invitrogen #A11030) were applied and incubated for 1 hour at room temperature to detect primary antibody binding. Slowfade gold was used as an antifade mounting medium.

Slides were imaged under a 25x water immersion objective using a Leica SP5 II Confocal microscope. All images were taken sequentially with a z-stack step size at 1 µm, a line average of 2, speed of 200 Hz, and a resolution of 1024 × 1024 pixels. Image J was used to compile slices into a maximum intensity projection(Jaeger et al., 2018).

### Statistical analysis

All statistical analyses were executed using GraphPad Prism 6 software. Sample size and statistical tests performed are provided in the Figure legends. For Dunnett’s post hoc analyses, the experimental group was compared to all controls and the highest p-value reported over the experimental bar. Replicates are biological replicates, using different individual flies from 2 or more crosses. Sample sizes were based on previous experiments in which effect size was determined. Data was excluded on the basis of STROBE technical malfunctions for individual flies and criteria for data exclusion are as follows: i) if the light system was not working during training for individual arenas ii) if during training or testing a fly did not meet a standard minimum # of interactions for that genotype iii) if during training or testing the STROBE recorded an abnormally large # of interactions for that genotype iiii) technical malfunctions due to high channel capacitance baseline activity v) if a fly was dead in an arena.

## DATA AVAILABILITY

All data are available in the manuscript and Supplementary Information, or are available upon request from the corresponding author.

## ACKNOWLEDGEMENTS

We thank Celia Lau for the original SEZ diagram models, and members of the Gordon lab for comments on the manuscript. This work was funded by Natural Sciences and Engineering Research Council (NSERC) grants RGPIN-2016–03857 and RGPAS 492846–16.

## AUTHOR CONTRIBUTIONS

M.J, P.Y.M, and M.D.G conceived the project. M.J and M.D.G wrote the manuscript. M.J performed all experiments, analyzed the data, and made the figures. P.Y.M and P.J offered experimental advice. M.D.G supervised the project.

## DECLARATION OF INTERESTS

The authors declare no competing interests.

## SUPPLEMENTAL INFORMATION

Supplemental information includes 3 figures.

## Supplemental Information

**Figure S1:**
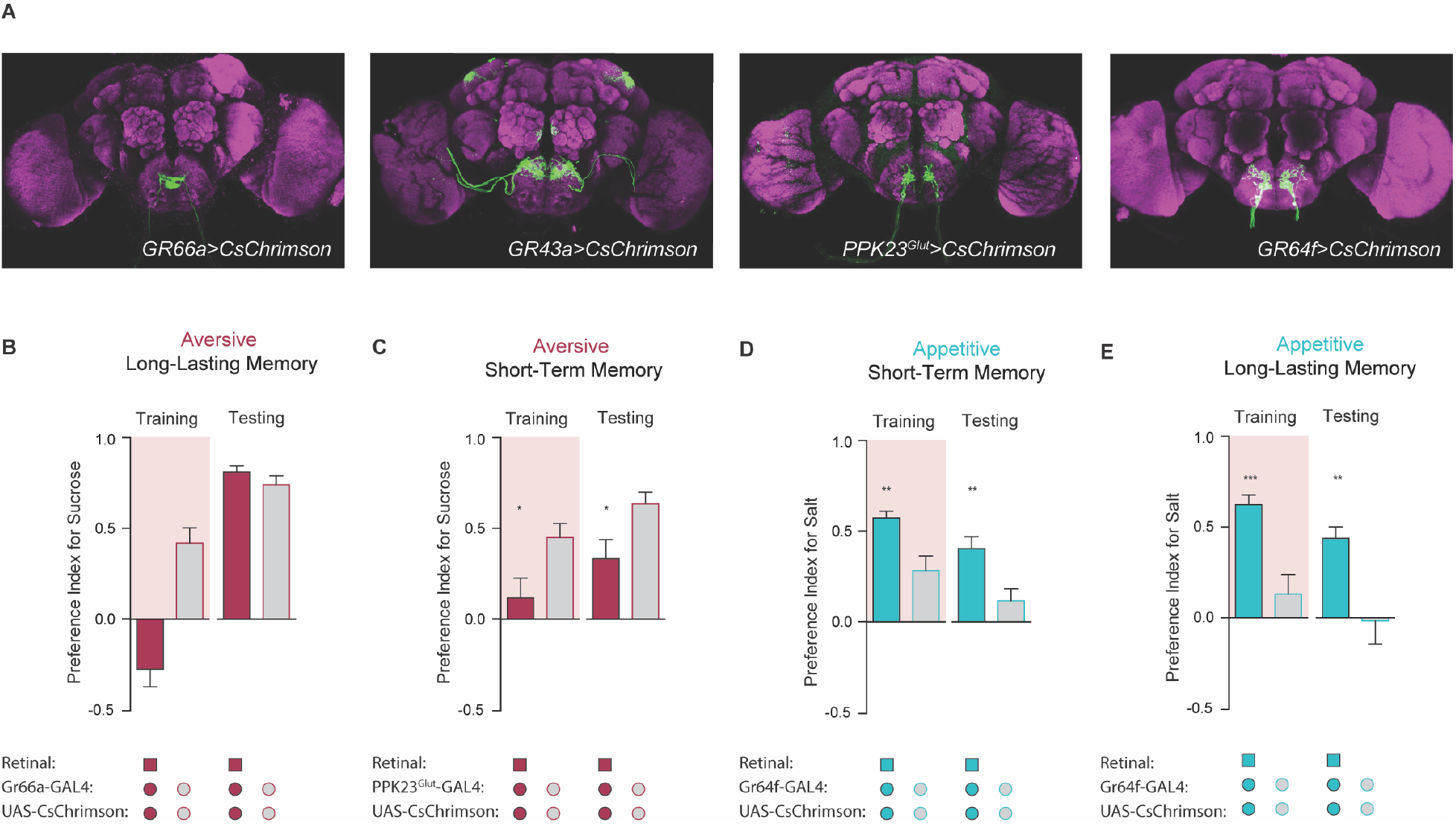
Gustatory receptor neuron activation produces reward and punishment signals in taste memory formation. (A) Projections from *GR66a, Gr43a, PPK23*^*Glut*^ *and GR64f* GRNs. (B) Preference index for *Gr66a>CsChrimson* flies fed all-*trans-*retinal in the long-lasting taste memory assay with or without retinal (n=19-21). (C) Preference indices for *PPK23*^*Glut*^*>CsChrimson* flies in the short-term taste memory assay with or without retinal (n=22-30). (D, E) Preference index for *Gr64f>CsChrimson* flies in the short-term (n=28-36) (D) and long-lasting (n=13-21) (E) taste memory assay with or without retinal. Preference indices are mean ± SEM, *t*-test: \**p* < 0.05, \*\**p* < 0.01, \*\*\**p* < 0.001.

**Figure S2:**
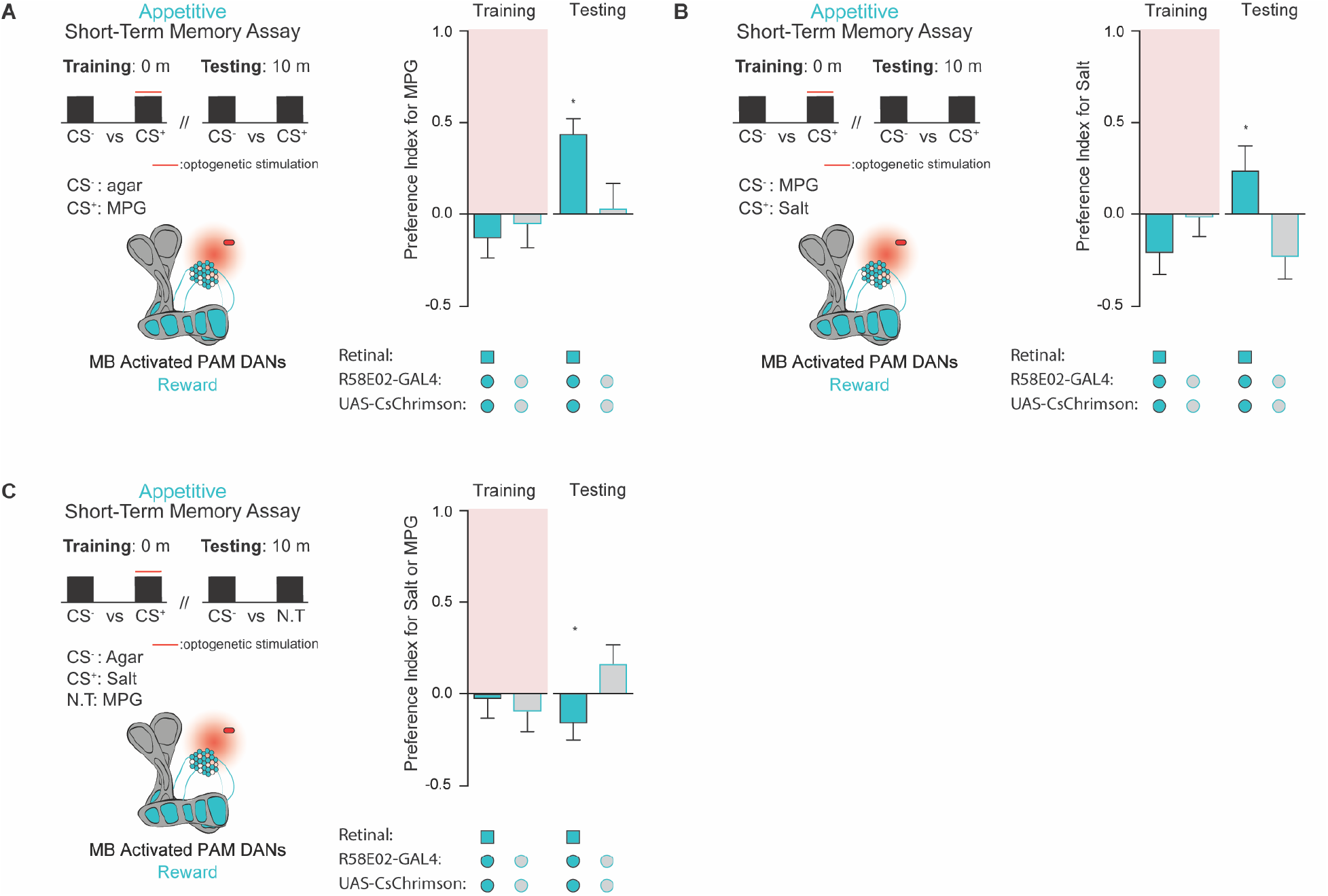
Taste memories are specific to the CS^+^. (A) Schematic outlining STM assay and MB compartments innervated by PAM driver R58E02. Preference index of *R58E02>CsChrimson* flies in the STM assay when monopotassium glutamate (MPG) is used as a CS^+^ (n=16-19). (B) Preference index of *R58E02>CsChrimson* flies in the STM assay with salt as a CS^+^ and MPG as a CS^-^ (n=16-19). (C) Preference indices of *R58E02>CsChrimson* flies in the short-term memory assay when the CS^+^ (NaCl) is switched for to the novel tastant (N.T.) MPG during testing (n=21-30). Preference indices are mean ± SEM, *t*-test, : \**p* < 0.05.

**Figure S3:**
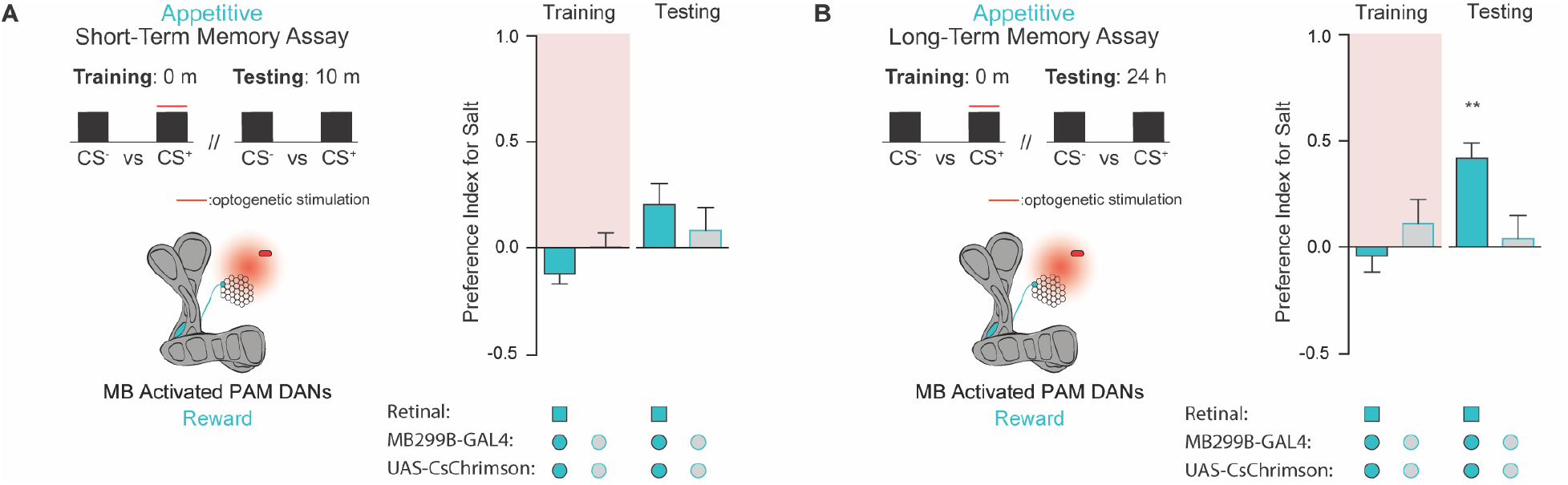
Activation of discrete PAM subpopulations induces distinct types of taste memories. (A, B) Schematic outlining STM and LTM assays, and an alternate PAM-α1 driver, innervating the α1 compartment of the horizontal MB lobe. Preference indices of *MB299B>CsChrimson* flies fed all-*trans-*retinal in the short-term (n=18-23) (A) and long-term (n=22-28) (B) memory assays compared to controls. Preference indices are mean ± SEM, *t*-test: \*\**p* < 0.01. (n=18-26)

## Notes

### Competing Interest Statement

The authors have declared no competing interest.

### Summary of Updates

We have added some additional control experiments to validate the paradigm, rewritten parts of the text, and made minor updates to the graphics in the figures.

