## Supplemental figures for "Optogenetic induction of appetitive and aversive taste memories in *Drosophila*"

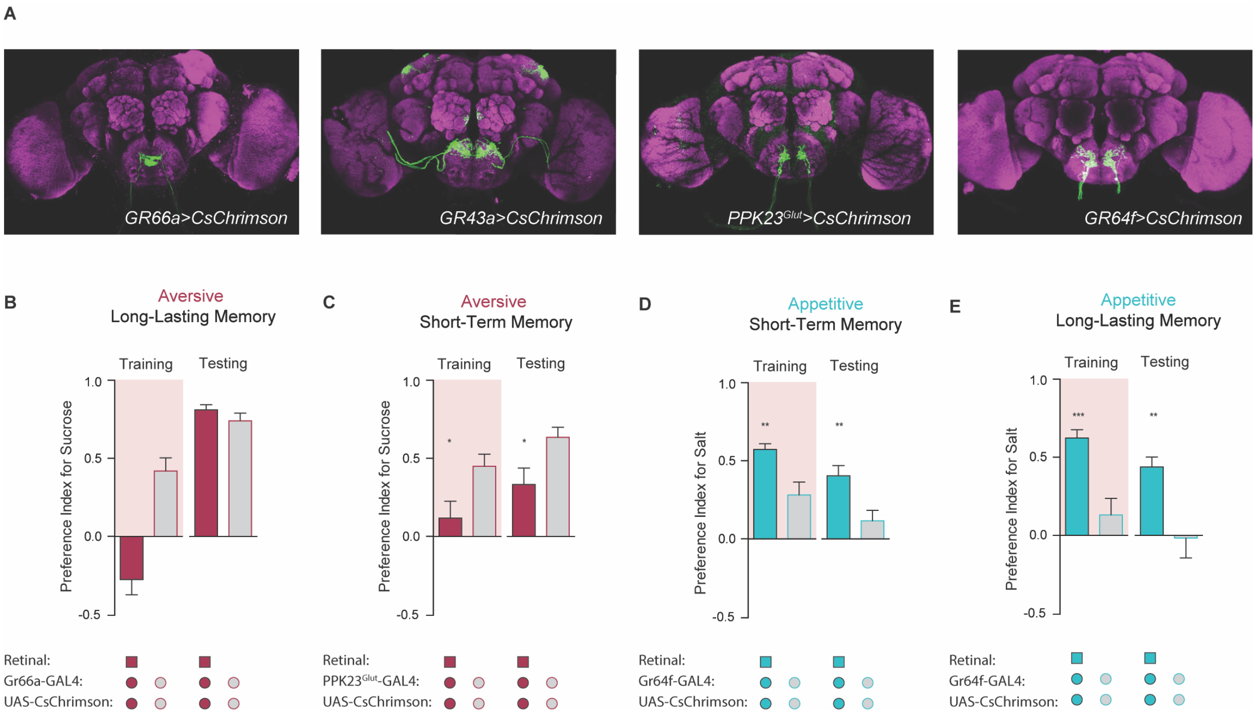


**Figure S1: Gustatory receptor neuron activation produces reward and punishment signals in taste memory formation.** (A) Projections from *GR66a*, *Gr43a*, *PPK23*$^{Glut}$ *and GR64f* GRNs. (B) Preference index for *Gr66a>CsChrimson* flies fed all-*trans*-retinal in the long-lasting taste memory assay with or without retinal (n=19-21). (C) Preference indices for *PPK23*$^{Glut}$*>CsChrimson* flies in the short-term taste memory assay with or without retinal (n=22-30). (D, E) Preference index for *Gr64f>CsChrimson* flies in the short-term (n=28-36) (D) and long-lasting (n=13-21) (E) taste memory assay with or without retinal. Preference indices are mean ± SEM, *t*-test: $*p < 0.05$, $**p < 0.01$, $***p < 0.001$.

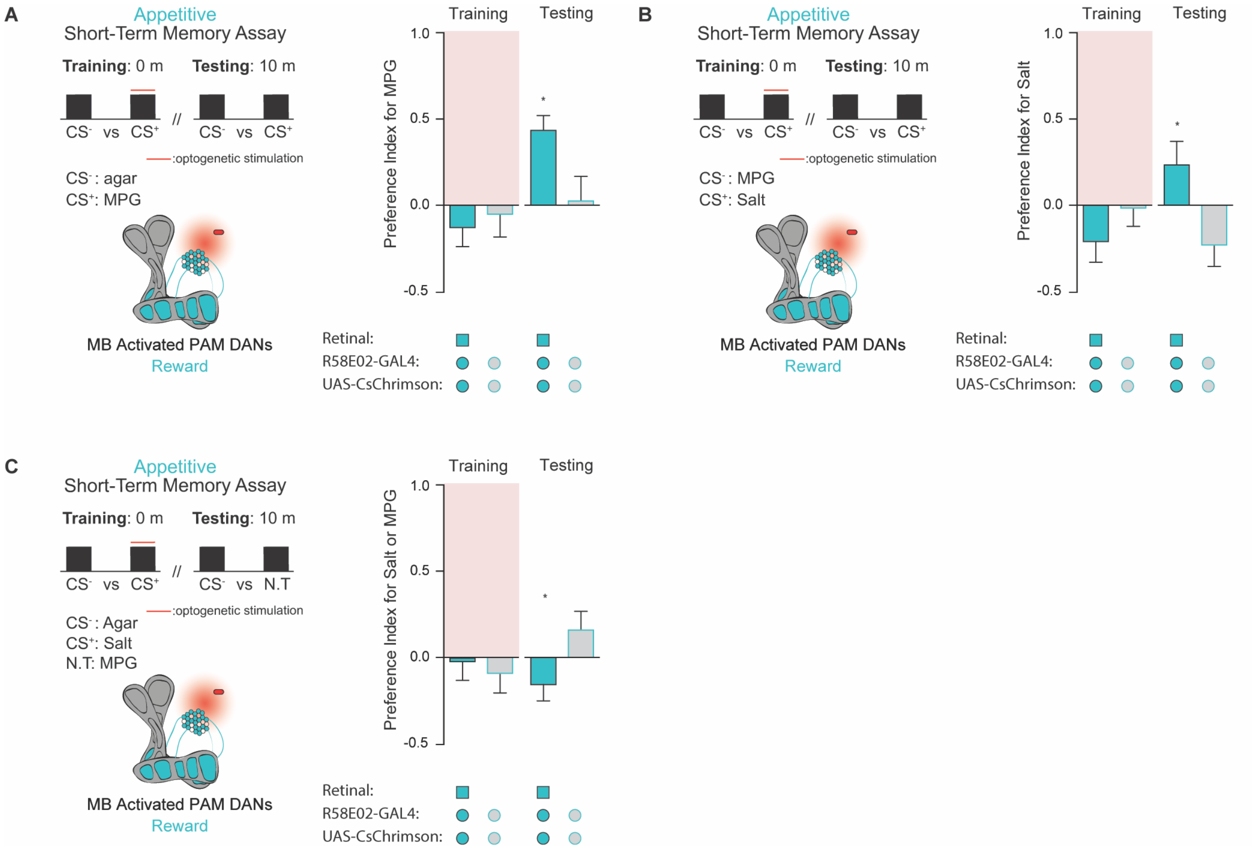


**Figure S2: Taste memories are specific to the CS+.** (A) Schematic outlining STM assay and MB compartments innervated by PAM driver R58E02. Preference index of *R58E02>CsChrimson* flies in the STM assay when monopotassium glutamate (MPG) is used as a CS+ (n=16-19). (B) Preference index of *R58E02>CsChrimson* flies in the STM assay with salt as a CS+ and MPG as a CS- (n=16-19). (C) Preference indices of *R58E02>CsChrimson* flies in the short-term memory assay when the CS+ (NaCl) is switched for to the novel tastant (N.T.) MPG during testing (n=21-30). Preference indices are mean ± SEM, *t*-test, : $*p < 0.05$.

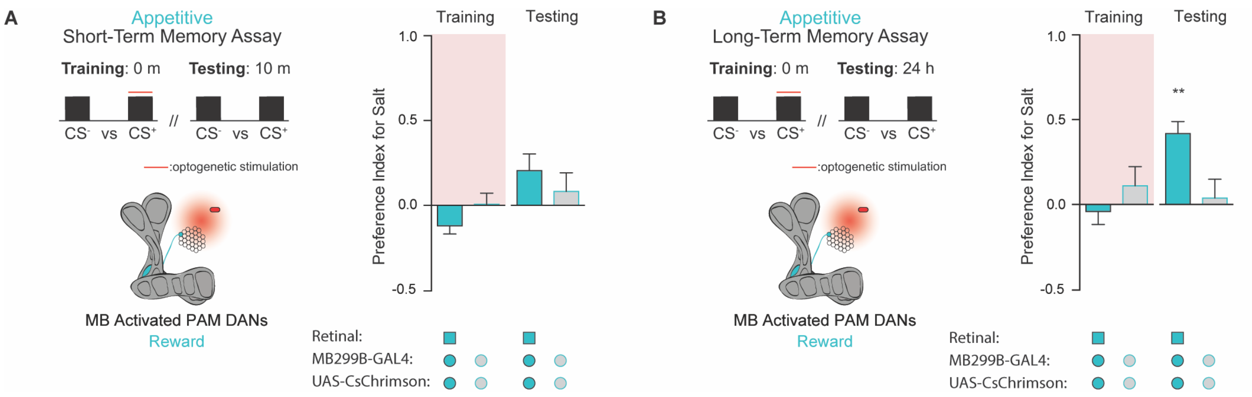


**Figure S3: Activation of discrete PAM subpopulations induces distinct types of taste memories.** (A, B) Schematic outlining STM and LTM assays, and an alternate PAM-α1 driver, innervating the α1 compartment of the horizontal MB lobe. Preference indices of *MB299B>CsChrimson* flies fed all-*trans*-retinal in the short-term (n=18-23) (A) and long-term (n=22-28) (B) memory assays compared to controls. Preference indices are mean ± SEM, *t*-test: $**p < 0.01$. (n=18-26)
